# Crystal structure of an RNA/DNA strand exchange junction

**DOI:** 10.1101/2022.01.24.477483

**Authors:** Joshua C. Cofsky, Gavin J. Knott, Christine L. Gee, Jennifer A. Doudna

## Abstract

Short segments of RNA displace one strand of a DNA duplex during diverse processes including transcription and CRISPR-mediated immunity and genome editing. These strand exchange events involve the intersection of two geometrically distinct helix types—an RNA:DNA hybrid (A-form) and a DNA:DNA homoduplex (B-form). Although previous evidence suggests that these two helices can stack on each other, it is unknown what local geometric adjustments could enable A-on-B stacking. Here we report the X-ray crystal structure of an RNA-5’/DNA-3’ strand exchange junction at an anisotropic resolution of 1.6 to 2.2 Å. The structure reveals that the A-to-B helical transition involves a combination of helical axis misalignment, helical axis tilting and compression of the DNA strand within the RNA:DNA helix, where nucleotides exhibit a mixture of A- and B-form geometry. These structural principles explain previous observations of conformational stability in RNA/DNA exchange junctions, enabling a nucleic acid architecture that is repeatedly populated during biological strand exchange events.

## Introduction

Although structural and mechanistic information is available for various types of DNA strand exchange processes [1–8], comparatively little is known about RNA/DNA strand exchange. In this reversible process, a strand of RNA hybridizes to one strand of a DNA duplex while displacing the other strand, requiring concomitant disruption of DNA:DNA base pairs and formation of RNA:DNA base pairs. This process occurs most notably at the boundaries of R-loops, such as those left by transcriptional machinery [9], those employed by certain transposons [10,11], or those created by CRISPR-Cas (<u>c</u>lustered <u>r</u>egularly <u>i</u>nterspaced <u>s</u>hort <u>p</u>alindromic <u>r</u>epeats, <u>C</u>RISPR-<u>as</u>sociated) enzymes during prokaryotic immunity or eukaryotic genome editing [12–15]. Structural insight into RNA/DNA strand exchange could therefore improve our understanding of how transcriptional R-loops are resolved and how CRISPR-Cas enzymes such as Cas9 manipulate R-loops to efficiently reject off-target DNA and recognize on-target DNA.

The defining feature of RNA/DNA strand exchange is the junction where the RNA:DNA helix abuts the DNA:DNA helix. Previous experiments on exchange junctions containing an RNA-5’ end and a DNA-3’ end (an “RNA-5’/DNA-3’ junction,” which is the polarity generated by Cas9) showed the component DNA:DNA duplex to be more thermodynamically stable than a free DNA helix end, perhaps due to interhelical RNA:DNA/DNA:DNA stacking [16]. While stacking in DNA-only junctions is thought to occur as it would in an uninterrupted B-form duplex [8,17,18], an analogous structural prediction cannot be made for RNA/DNA junctions because the two component helices are predisposed to different geometries: B-form for the DNA:DNA helix and a variant of A-form for the RNA:DNA helix [19–21]. A conformation that preserves base stacking across such a junction must reconcile base pairs that are flat and centered (B-form) with base pairs that are inclined and displaced from the helical axis (A-form). While prior structural studies of Okazaki fragments reckoned with a similar geometric puzzle [22], Okazaki fragments bear an RNA-3’/DNA-5’ polarity (opposite of the polarity addressed here) and lack the strand discontinuity that defines exchange junctions. Thus, the structural basis for the putative stacking-based stability in RNA-5’/DNA-3’ junctions remains unknown.

Here we present the X-ray crystal structure of an RNA-5’/DNA-3’ strand exchange junction, which undergoes an A-to-B transition without loss of base pairing or stacking across the exchange point. This structure reveals the principles of global helical positioning and local adjustments in nucleotide conformation that allow RNA:DNA duplexes to stack on DNA:DNA duplexes in the RNA-5’/DNA-3’ polarity. This model also complements previously determined cryo-electron microscopy structures of DNA-bound Cas9 for which poor local resolution in the original maps prevented accurate modeling of the leading R-loop edge.

## Results

Inspired by previous crystallographic studies of double-stranded DNA dodecamers [23,24], we designed crystallization constructs that contained a “template” DNA strand (12 nucleotides) and two “exchanging” RNA and DNA oligonucleotides that were complementary to each half of the template DNA strand. In different versions of these constructs, we varied the polarity (RNA-5’/DNA-3’ vs. RNA-3’/DNA-5’) and the internal termini, which were either flush (exchanging oligonucleotides were 6-mers) or extended with a one-nucleotide flap that was not complementary to the template strand (exchanging oligonucleotides were 7-mers, “flapped”). Only the flapped construct in the RNA-5’/DNA-3’ polarity (Fig 1A) yielded well-diffracting crystals (anisotropic resolution of 1.6 to 2.2 Å). Thus, all results discussed here describe a flapped RNA-5’/DNA-3’ strand exchange junction, which is the polarity previously observed to stabilize the component DNA:DNA duplex [16].

**Fig 1.**
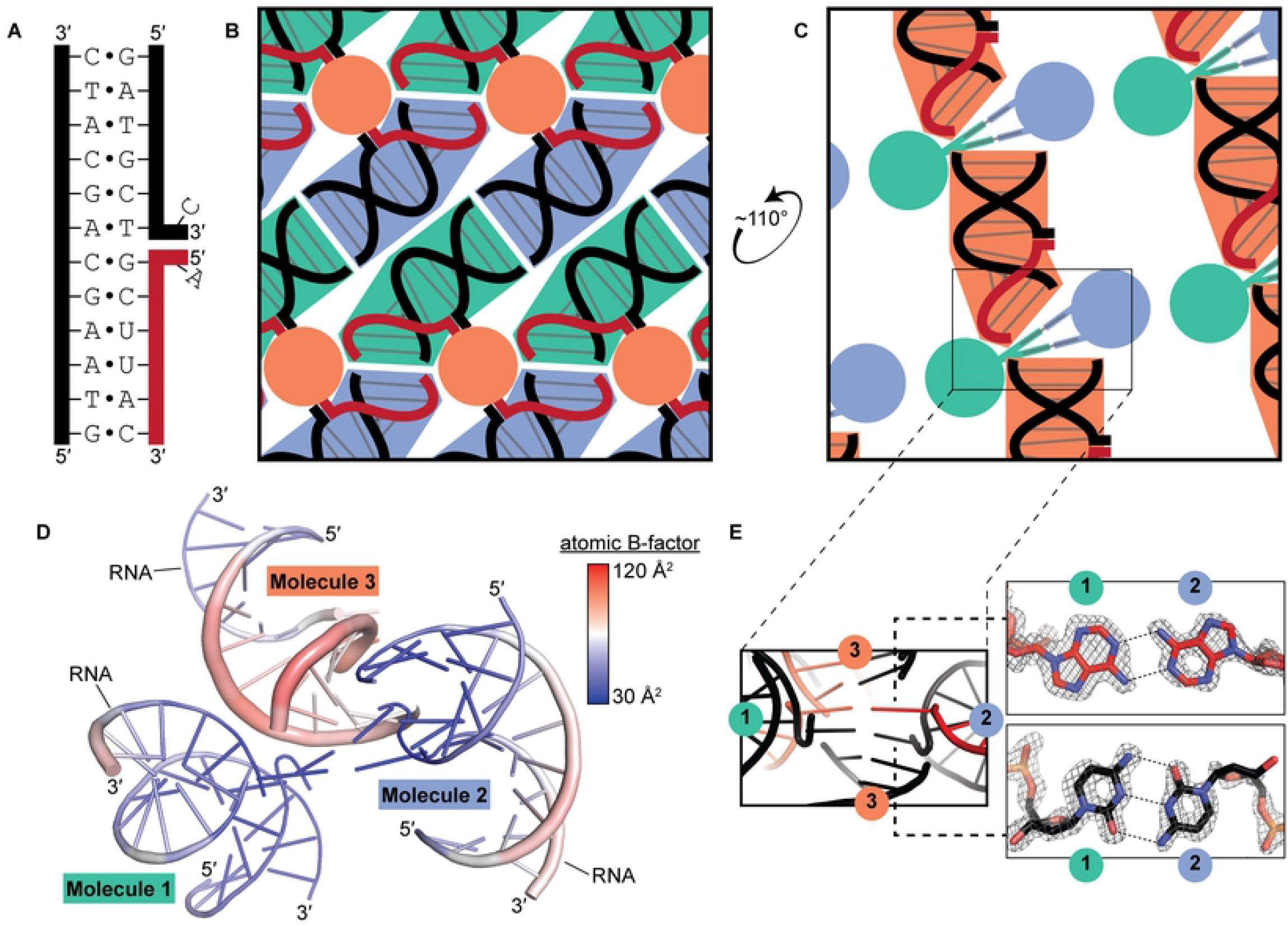
Stabilizing features of the crystal lattice. (A) Crystallization construct sequence. Black, DNA; red, RNA. (B) Schematized drawing (not to scale) of the crystal lattice along a direction that depicts the helical network formed by Molecules 1 and 2. Green shading, Molecule 1; blue shading, Molecule 2; orange shading, Molecule 3 (cross section). (C) Similar to panel B, but along a direction that depicts the helical network formed by Molecule 3. (D) Asymmetric unit colored by atomic B-factor. The thickness of the cartoon model also reflects the local B-factors. (E) Model and 2mF_o_-DF_c_ map (sharpened by −38 Å^2^, displayed at 3.3σ) of the Ade-Ade and Cyt-Cyt base pairs (contributed by the flap nucleotides of Molecules 1 and 2) that bridge the helical network formed by Molecule 3. Distortion in the map is due to diffraction anisotropy (see Methods).

We determined the X-ray crystal structure of the exchange junction (Table 1). In this structure, the asymmetric unit contains three molecules (a “molecule” comprises one DNA 12-mer and its complementary RNA and DNA 7-mers). The crystal lattice is largely stabilized by nucleobase stacking interactions both within and between molecules. Along one lattice direction, Molecules 1 and 2 form a continuous network of stacked helices, in which the external RNA:DNA duplex terminus of each Molecule 1 stacks on the equivalent terminus of Molecule 2, with a similar reciprocal interaction for the external DNA:DNA duplex termini (a “head-to-head” and “foot-to-foot” arrangement) (Fig 1B). Along another lattice direction, symmetry-related instances of Molecule 3 create a head-to-foot helical network (Fig 1C). Compared to Molecules 1 and 2, Molecule 3 is poorly ordered (Fig 1D), and its atomic coordinates appear less constrained by the data due to diffraction anisotropy (see Methods). In the Molecule 3 helical network, two base pairs formed between the flapped nucleotides of Molecules 1 and 2 bridge the duplex ends. The bridging nucleotides form a type I adenine-adenine (ribonucleotide) base pair and a type XV hemiprotonated cytosine-cytosine (deoxyribonucleotide) base pair [25] (Fig 1C, E).

**Table 1.** Crystallographic data and refinement statistics.

|  | RNA-5'/DNA-3' strand exchange junction (PDB XXXX) |
| --- | --- |
| **Data collection** |  |
| Wavelength (Å) | 1.116 |
| Resolution range (Å) | 35.335 - 1.637 (1.781-1.637) |
| Diffraction limit #1 (Å) | 1.656 |
| Principal axes (orthogonal basis) | 0.8648, -0.0396, -0.5006 |
| Principal axes (reciprocal lattice) | 0.657 a* - 0.168 b* - 0.735 c* |
| Diffraction limit #2 (Å) | 2.176 |
| Principal axes (orthogonal basis) | 0.1677, 0.9624, 0.2136 |
| Principal axes (reciprocal lattice) | 0.152 a* + 0.981 b* + 0.117 c* |
| Diffraction limit #3 (Å) | 1.637 |
| Principal axes (orthogonal basis) | 0.4733, -0.2686, 0.8389 |
| Principal axes (reciprocal lattice) | 0.397 a* - 0.342 b* + 0.852 c* |
| Space group | P 1 |
| Unit cell |  |
| a, b, c (Å) | 37.025, 43.586, 52.182 |
| $\alpha, \beta, \gamma$ (°) | 92.06, 103.72, 99.95 |
| Total reflections | 147975 (7830) |
| Unique reflections | 24808 (1240) |
| Multiplicity | 6.0 (6.3) |
| Spherical completeness (%) | 64.7 (14.5) |
| Ellipsoidal completeness (%) | 86.8 (50.0) |
| $\langle I/\sigma(I) \rangle$ | 15.6 (1.5) |
| Wilson B-factor (Å<sup>2</sup>) |  |
| Eigenvalue #1 (Å) | 48.57 |
| Principal axes (orthogonal basis) | 0.9603, -0.1659, -0.2243 |
| Principal axes (reciprocal lattice) | 0.799 a* - 0.323 b* - 0.507 c* |
| Eigenvalue #2 (Å) | 86.73 |
| Principal axes (orthogonal basis) | 0.2257, 0.9345, 0.2753 |
| Principal axes (reciprocal lattice) | 0.209 a* + 0.961 b* + 0.183 c* |
| Eigenvalue #3 (Å) | 45.47 |
| Principal axes (orthogonal basis) | 0.1639, -0.3150, 0.9348 |
| Principal axes (reciprocal lattice) | 0.123 a* - 0.300 b* + 0.946 c* |
| R<sub>merge</sub> | 0.037 (1.293) |
| R<sub>meas</sub> | 0.041 (1.410) |
| R<sub>pim</sub> | 0.016 (0.556) |
| CC<sub>1/2</sub> | 0.999 (0.474) |
| **Refinement** |  |
| Resolution range (Å) | 35.34 - 1.641 (1.768 - 1.641) |
| Reflections used in refinement | 24717 (1054) |
| Reflections used for R<sub>free</sub> | 1223 (41) |
| R<sub>work</sub> | 0.2371 (0.3692) |
| R<sub>free</sub> | 0.2846 (0.3561) |
| CC<sub>work</sub> | 0.912 (0.616) |
| CC<sub>free</sub> | 0.939 (0.562) |
| Number of non-hydrogen atoms | 1651 |
| macromolecules | 1584 |
| ligands | 0 |
| solvent | 67 |
| Protein residues | 0 |
| RMSD – bond lengths (Å) | 0.014 |
| RMSD – angles (°) | 1.42 |
| Clashscore | 0.00 |
| Average B-factor | 59.75 |
| macromolecules | 60.15 |
| solvent | 50.35 |
| Number of TLS groups | 15 |

Diffraction limits and eigenvalues of overall anisotropy tensor on $|F|$ s are displayed alongside the corresponding principal axes of the ellipsoid fitted to the diffraction cut-off surface as direction cosines in the orthogonal basis (standard PDB convention), and in
terms of reciprocal unit-cell vectors. Statistics for the highest-resolution shell are shown in parentheses.

The three molecules of the asymmetric unit exhibit canonical Watson-Crick base pairing at all twelve nucleotides of the template DNA strand, and they are generally similar in conformation (RMSD_Mol1,mol2_=0.703 Å; RMSD_MOl1,MOl3_=1.528 Å, RMSD_MOl2,MOl3_=1.770 Å) (Fig 2A). The most dramatic differences are between Molecules 1/2 and Molecule 3. For example, Molecule 3’s flapped nucleotides form no intermolecular base pairs, and the conformation of the DNA flap is flipped relative to Molecules 1/2. Additionally, the external three base pairs of Molecule 3’s DNA:DNA helix tilt slightly toward the major groove as compared to the equivalent positions of Molecules 1/2. Notably, the similarity of all three molecules at the three base pairs on either side of the exchange point (RMSD_MOl1,MOl2_=0.567 Å; RMSD_MOl1,MOl3_=0.502 Å, RMSD_Mol2,Mol3_=0.751 Å) suggests that the conformation in this region represents a low-energy solution to the stacking of RNA:DNA and DNA:DNA helices.

**Fig 2.**
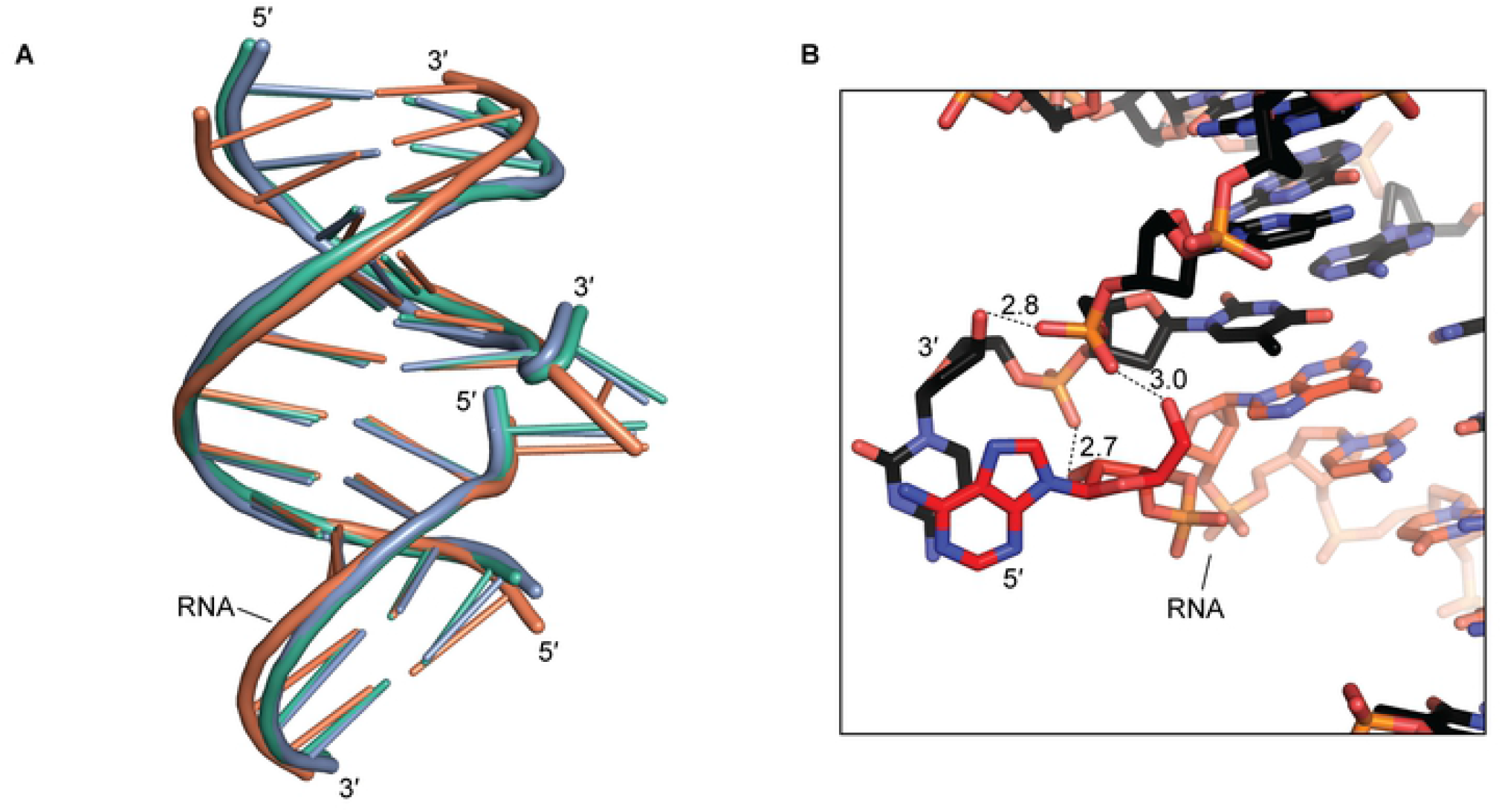
Molecule-to-molecule similarity and hydrogen bonding at the flapped nucleotides. (A) All-atom alignment of the three molecules in the asymmetric unit. Green, Molecule 1; blue, Molecule 2; orange, Molecule 3. Molecules 2 and 3 were aligned to Molecule 1 in this depiction. (B) Hydrogen bonding at the flapped nucleotides of Molecule 1. Dotted lines indicate hydrogen bonds, and adjacent numbers indicate interatomic distance in Å. Black, DNA; red, RNA. This hydrogen bonding pattern is also observed in Molecule 2 but not in Molecule 3.

At the exchange point of Molecules 1 and 2, the flapped nucleotides are stabilized not only by intermolecular base pairing (Fig 1C, E) and intramolecular stacking (Fig 2B), but also by hydrogen bonds between sugar hydroxyls and backbone phosphates. Specifically, at the junction-proximal phosphodiester within the DNA:DNA helix, the *pro-S*_p_ and *pro-R*_p_ oxygens are hydrogen-bonded to the terminal 3’ hydroxyl of the flapped DNA nucleotide and the terminal 5’ hydroxyl of the flapped RNA nucleotide, respectively. Additionally, the *pro-S*_p_ oxygen of the flapped DNA nucleotide is hydrogen-bonded to the 2’ hydroxyl of the flapped RNA nucleotide (Fig 2B). If the flaps were longer than one nucleotide, as would occur during biological strand exchange events, the hydrogen bonds to the terminal 3’/5’ hydroxyls would be perturbed. However, in Molecule 3, the flipped deoxycytidine conformation precludes all the mentioned extrahelical hydrogen bonds, yet the base-paired nucleotides within the junction are conformationally similar to the same region in Molecules 1 and 2 (Fig 2A). Therefore, we expect that the structural features of interest to this work—that is, the conformation of the base-paired nucleotides immediately adjacent to the junction—would be populated by junctions bearing flush RNA/DNA ends or flaps of arbitrary length.

To understand the nature of the transition in helical geometry across the junction, we performed alignments of regularized A-form and B-form DNA:DNA helices with the observed RNA:DNA and DNA:DNA helices, respectively. These alignments revealed that the DNA:DNA helix closely approximates perfect B-form geometry, especially in the nucleotides closest to the junction (Fig 3A-C). Likewise, the RNA strand of the RNA:DNA helix closely approximates A-form geometry (Fig 3A-C). On the other hand, the DNA strand of the RNA:DNA helix deviates from its A-form trajectory in the three nucleotides that approach the exchange point, where the backbone is compressed toward the minor groove (Fig 3B, D).

**Fig 3.**
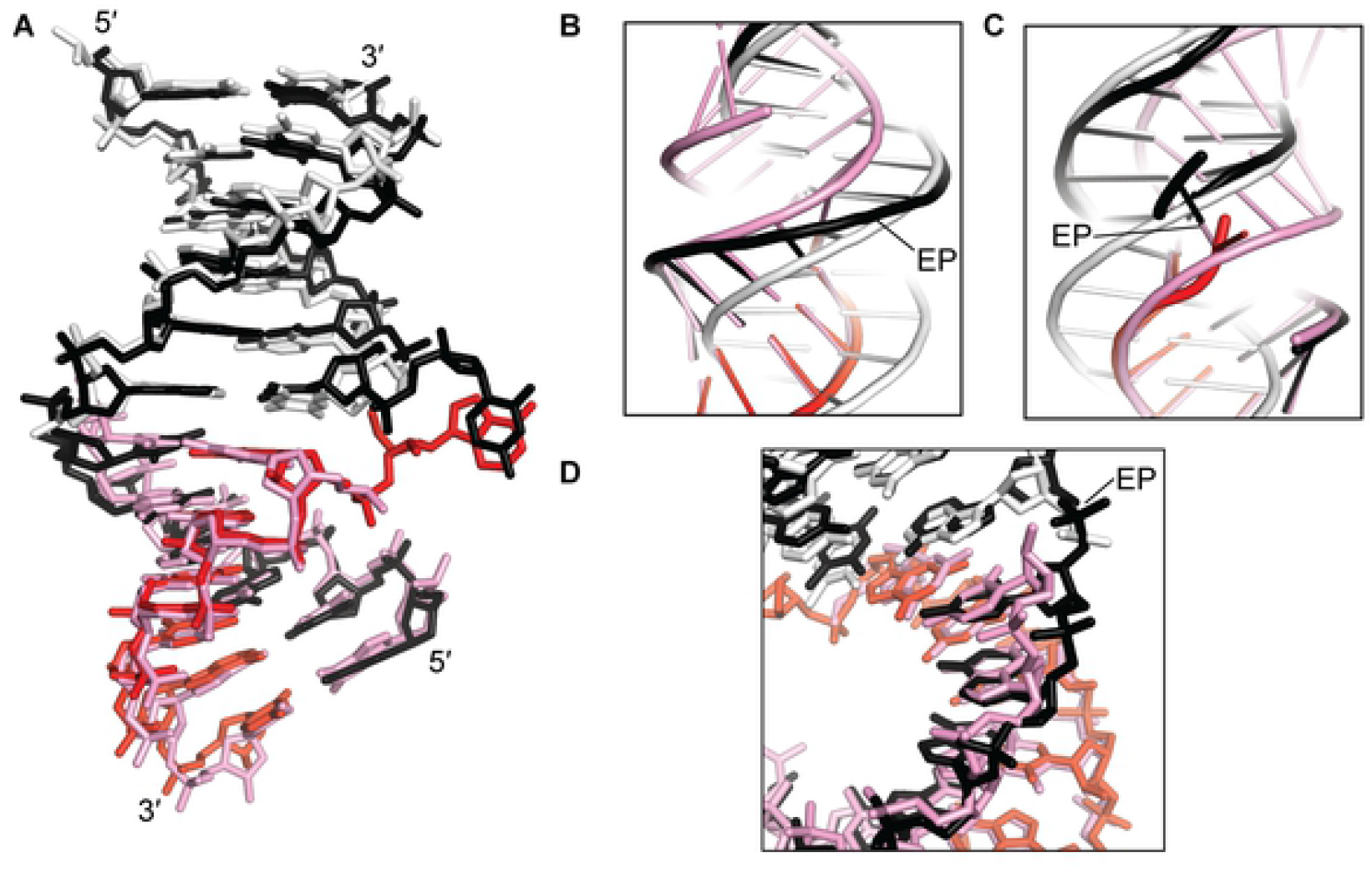
Alignments to regularized A-form/B-form helices. (A) Black, DNA of Molecule 1; red, RNA of Molecule 1; white, regularized B-form DNA:DNA helix aligned to the 6 bp of Molecule 1’s DNA:DNA helix; pink, regularized A-form DNA:DNA helix aligned to the 6 bp of Molecule 1’s RNA:DNA helix. (B) Cartoon depiction, focused on the continuous strand. The alignment procedure for each 6-bp block was identical to that performed in panel A, but in this depiction, the B-form (white) and A-form (pink) helices were extended by an additional 6 bp (extended nucleotides were not considered during alignment) to illustrate the path that the helix would take if continuing along a perfect B-form or A-form trajectory. EP, exchange point (that is, the phosphodiester or gap lying between the two nucleotides where the helix changes from RNA:DNA to DNA:DNA). (C) Similar to panel B, but focused on the discontinuous strand. (D) Close-up of the same representation depicted in panel A, focused on the nucleotides that deviate most dramatically from the aligned A-form helix.

Interestingly, calculation of *z_P_*, a geometric parameter that differentiates A-form from B-form base steps [26], indicated that the RNA:DNA base step adjacent to the exchange point is A-like, while the base steps in the center of the RNA:DNA helix are intermediate in their A/B character (Fig 4A). This result indicates an important distinction between strand trajectory (in terms of global alignment to a regularized A-form or B-form helix) and the local nucleotide conformations that underlie the trajectory. In the RNA:DNA helix, the departure from A-form trajectory observed at junction-adjacent nucleotides appears to result from non-A conformations at more junction-distal nucleotides. Other indicators of helical geometry also suggest a mixture of A and B character across the RNA:DNA helix (S1 Fig).

**Fig 4.**
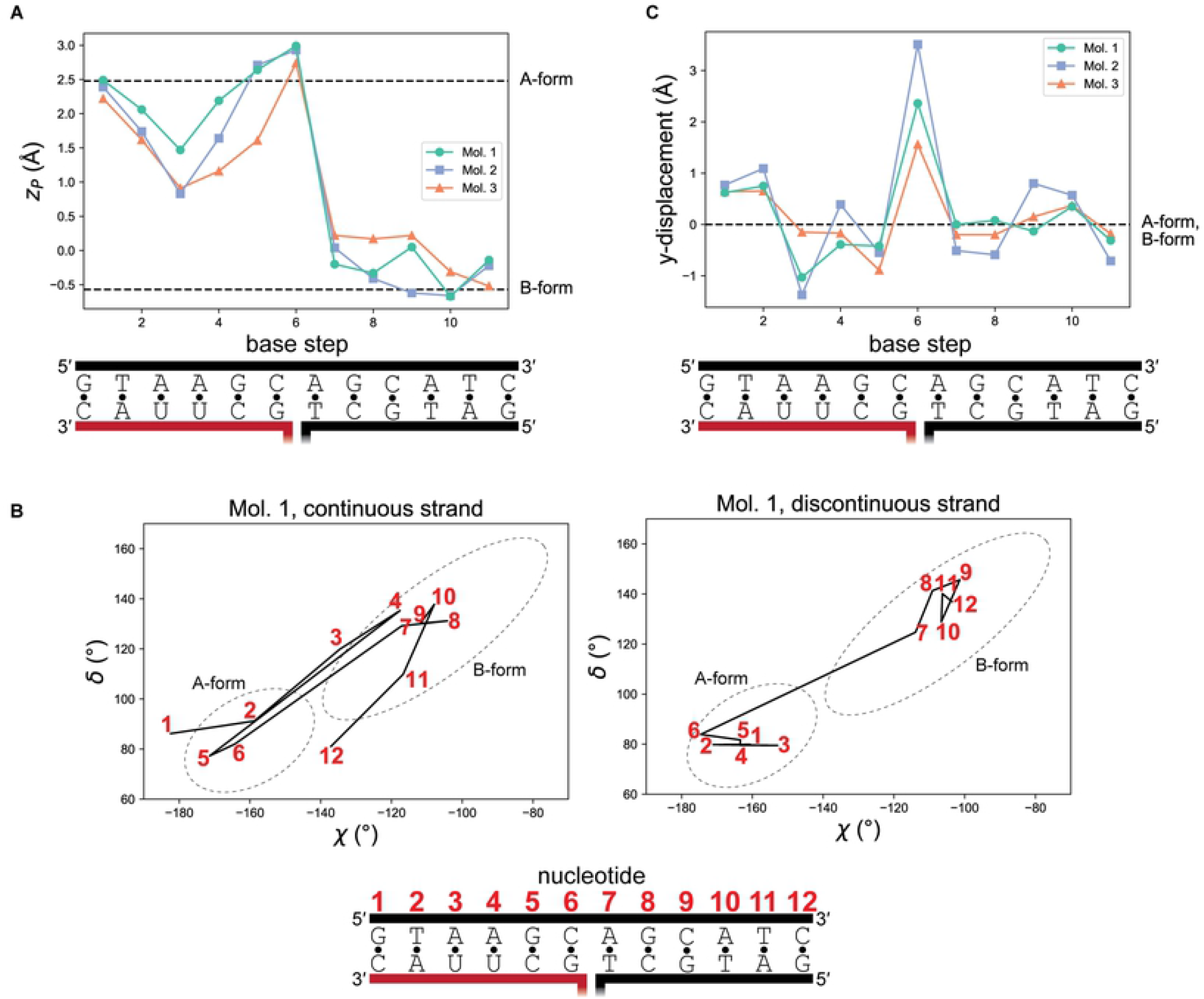
Geometric details of the A-to-B transition. (A) For a given base step, the parameter *z_P_* is the mean of the *z*-displacement of the two phosphorus atoms from the dimer’s reference *xy*-plane. Note that *z_P_* is defined by a pair of dinucleotides, so there are only 11 data points for a 12-bp helix, and integral x-values lie between the base pairs in the diagram. This parameter was originally introduced for its utility in distinguishing A-form from B-form base steps. Black, DNA; red, RNA. (B) χ and δ are the two nucleotide torsion angles that best distinguish A-form from B-form geometry. Note that these torsion angles are defined for each individual nucleotide, so there are 24 data points for a 12-bp helix. Integers in red refer to individual nucleotides, as indicated in the schematic at the bottom. Dashed ellipses were drawn to match those depicted in [27]. (C) Y-displacement. Similar to *z_P_*, this parameter describes base steps (pairs of dinucleotides), not individual nucleotides. This parameter cannot distinguish A-form from B-form geometry. Instead, note that the base step across the exchange point dramatically departs from both A-form and B-form geometry.

To probe helical geometry with strand specificity, we calculated χ and δ, nucleotide torsion angles that differ in A-form vs. B-form helices [27]. These parameters revealed that the irregularities observed in the paired base step parameters (Fig 4A and S1 Fig) arise entirely from the template DNA strand, which flips between A- and B-like conformations within the RNA:DNA hybrid (Fig 4B and S2 Fig). In contrast, the RNA strand is entirely A-like, and all nucleotides of the DNA:DNA helix are B-like except at position 12 of the continuous strand, which is likely due to an end effect. These observations agree with the conclusions drawn from the alignments (Fig 3A), and they highlight the DNA strand of the RNA:DNA helix as the structure’s most geometrically irregular region, which may enable the junction-adjacent deviation in trajectory.

In addition to the distortions in the continuous DNA strand, the geometric switch also seems to depend on the break in the discontinuous strand, which facilitates a marked jump in the backbone trajectory across the exchange point (Fig 3C). This feature reflects a global jump in helical positioning that is visualized most clearly in the aligned regularized A-form and B-form duplexes, whose helical axes are tilted and misaligned with respect to each other (the helical axes are tilted from parallel by 14°, Mol1; 18°, Mol2; 2°, Mol3) (Fig 2A and Fig 3B, C). Axis misalignment is detectable in the large positive y-displacement value across the central base step, which deviates dramatically from the expected value (0 Å) for either an A-form or B-form duplex (Fig 4C). This observation emphasizes the exchange point as a special base step with noncanonical alignment, made possible by discontinuity in the exchanging strands.

## Discussion

Together, our data suggest that stacking an RNA:DNA helix on a DNA:DNA helix does not require deviation of the RNA strand or either strand of the DNA:DNA helix from their native A-form or B-form conformations, respectively. Instead, continuous stacking appears to result from a combination of three structural principles. First, alternating A-like and B-like nucleotide conformations in the hybrid’s DNA strand compress the strand relative to a pure A-form trajectory (Fig 3B, D, Fig 4B, Fig 5A). Due to A-form base pair inclination (~20° from perpendicular to the helical axis) in RNA:DNA duplexes, the DNA naturally juts further along the helical axis than the RNA at the RNA-5’ end. This slanted RNA:DNA end can be stacked upon a flat DNA:DNA end through strand-specific compression—that is, compression of the hybrid’s protruding DNA strand (Fig 5A). Second, an alternative to strand compression is to tilt the helical axes themselves, which occurs in Molecules 1 and 2 but not Molecule 3 (Fig 2A and Fig 5A). Third, the helical centers are misaligned at the exchange point (Fig 3B, C and Fig 4C), which effectively aligns the off-center base pairs of the A-form duplex with the centered base pairs of the B-form duplex (Fig 5B).

**Fig 5.**
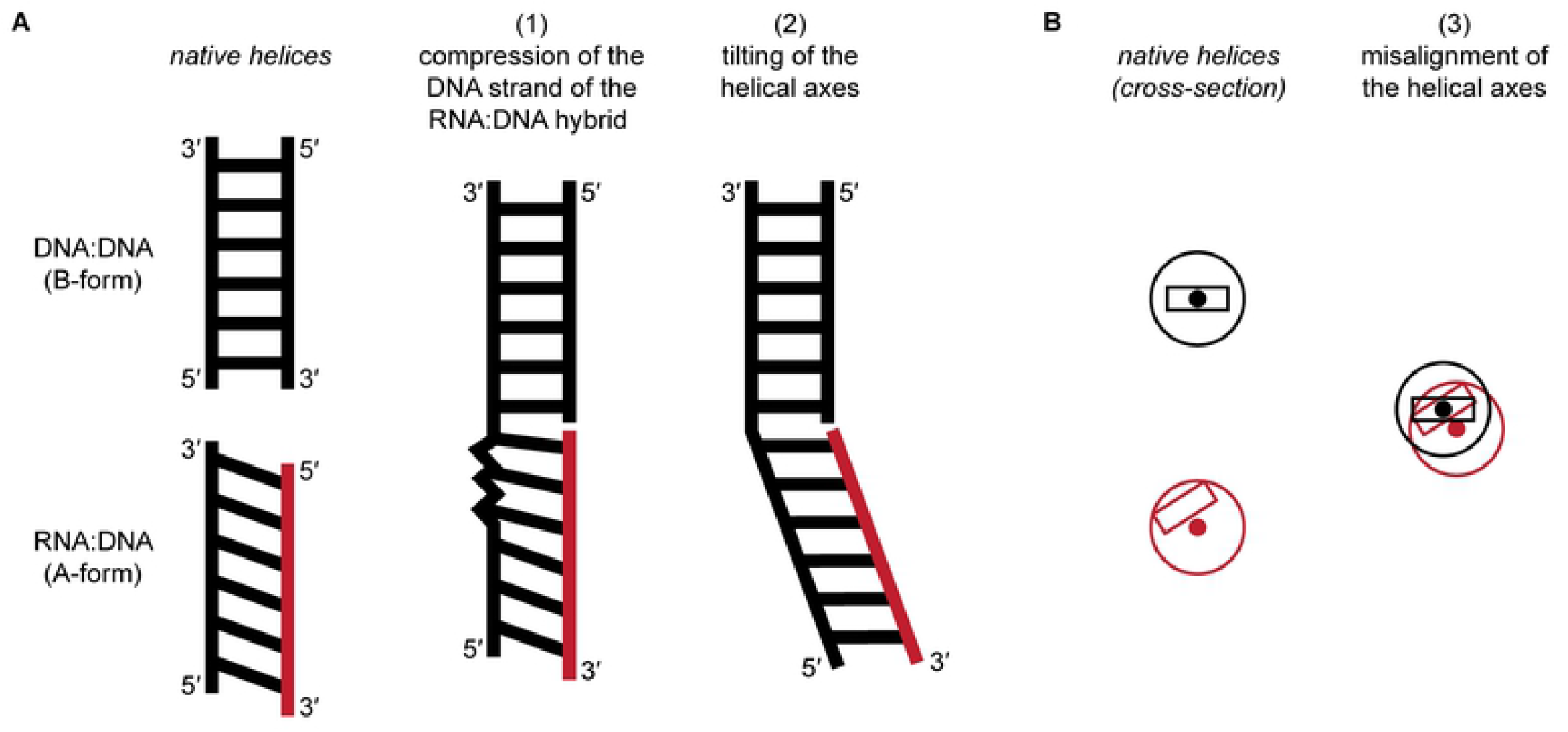
Structural principles of A-on-B stacking at the RNA-5’/DNA-3’ strand exchange junction. (A) Simplified schematics illustrating strand-specific compression and tilting of the helical axes. The slanted appearance of the RNA:DNA duplex is intended to represent the base pair inclination characteristic of A-form duplexes, which pushes the 3’ DNA end farther along the helical axis than the 5’ RNA end. Black, DNA; red, RNA. (B) Helical cross-sections. Black, DNA:DNA helix; red, RNA:DNA helix. The rectangle represents the base pair nearest the exchange point (centered in the B-form helix, off-center in the A-form helix). The solid circle represents the helical axis. The true stacking solution is a combination of the three principles illustrated here, although Molecule 3 does not exhibit tilting.

This new structure is best examined in the context of previous structural studies of RNA:DNA/DNA:DNA junctions emulating Okazaki fragments, which include a chimeric (covalently continuous) RNA-DNA strand. When crystallized, these fragments assumed an entirely A-form conformation, even within the DNA:DNA duplex [28–32]. However, in solution, Okazaki fragments resembled the present structure in that they were A-like within the RNA:DNA helix and B-like within the DNA:DNA helix [22,33–36]. Solution structures also exhibited a tilt between the RNA:DNA/DNA:DNA helical axes and intermediate nucleotide geometry within the DNA of the hybrid. Because intermediate geometry is a known feature of the DNA of any RNA:DNA hybrid [19,20], it may be the natural inclination of this more geometrically ambiguous strand to accommodate the A-to-B transition as it does in the present structure. Notably, dramatic misalignment of the RNA:DNA/DNA:DNA helical centers is observed only in the present structure and is likely enabled by the break in the exchanging strands, which is not a feature of Okazaki fragments.

Because stable stacking of another duplex on a DNA:DNA terminus is expected to inhibit duplex melting [37], the structural principles illuminated here may explain the rigidity that we previously observed in the DNA:DNA duplex of RNA-5’/DNA-3’ exchange junctions [16]. However, it is also possible that different sequences or environments promote different conformational preferences than those observed in this crystal structure. Previously, we also observed that the DNA:DNA duplex in junctions of the opposite polarity (RNA-3’/DNA-5’) is destabilized relative to a non-exchanging terminus [16]. Unfortunately, because that junction type failed to crystallize under our tested conditions, this odd asymmetry in junction structure remains unexplained.

Nevertheless, the stacked RNA-5’/DNA-3’ structure determined here represents a key conformation that is likely populated throughout RNA/DNA exchange events, including those mediated by the genome-editing protein Cas9. Branch migration is crucial to Cas9 target search, which involves repeated R-loop formation (RNA invades a DNA:DNA duplex) and resolution (DNA invades an RNA:DNA duplex) until the true target is located [15]. During this process, the leading R-loop edge likely passes through interhelically stacked states between base pair formation and breakage events. Consistent with this prediction, in some cryo-electron microscopy structures depicting Cas9-bound R-loops, the leading (RNA-5’/DNA-3’) R-loop edge appeared interhelically stacked [38,39]. While local resolution was insufficient to enable accurate atomic modeling of the exchange junction from the original electron microscopy maps, our high-resolution crystal structure provides a new geometric standard for modeling this kind of junction.

Importantly, exchange junctions are dynamic structures, and each time an R-loop grows or shrinks, stacking must be disrupted at the junction [8]. Thus, in addition to the stacked structure determined here, which can be interpreted as a ground state, strand exchange also requires passage through unstacked conformations, some of which may resemble the junction structures seen in other Cas9-bound R-loops [40,41]. A complete model of RNA/DNA strand exchange, then, will rely on a structural and energetic understanding of the junction in both stacked and unstacked states, and it will account for the effects of the proteins acting in R-loop formation and resolution.

## Methods

### Oligonucleotide synthesis and sample preparation

All oligonucleotides (DNA 12-mer {5’-GTAAGCAGCATC-3’}; DNA 7-mer {5’-GATGCTC-3’}; RNA 7-mer {5’-AGCUUAC-3’}) were synthesized and purified by Integrated DNA Technologies (high-performance liquid chromatography (HPLC) purification for DNA oligonucleotides and RNase-free HPLC purification for the RNA oligonucleotide). Dry oligonucleotides were dissolved in nuclease-free water (Qiagen), and concentrations were estimated by Nanodrop (Thermo Scientific) absorbance measurements with extinction coefficients estimated according to [42] (DNA 12-mer, ε_260_=135200 M^-1^·cm^-1^; DNA 7-mer, ε_260_=70740 M^-1^·cm^-1^; RNA 7-mer, ε_260_=75580 M^-1^·cm^-1^). The three oligonucleotides were combined and diluted in water, each at 500 μM final concentration. This exchange junction sample was incubated at 50°C for 10 minutes, cooled to 25°C within a few seconds, and used directly in the crystallization setups described below.

### Crystallization and data collection

Initial screens were performed using Nucleix and Protein Complex suites (Qiagen) in a sitting-drop setup, with 200 nL of sample added to 200 nL of reservoir solution by a Mosquito instrument (SPT Labtech) and incubated at either 4°C or 20°C. Several conditions yielded crystals within one day, and initial hits were further optimized at a larger scale. The crystal used for the final dataset was produced as follows: 0.5 μL of sample was combined with 0.5 μL reservoir solution (0.05 M sodium succinate (pH 5.3), 0.5 mM spermine, 20 mM magnesium chloride, 2.6 M ammonium sulfate) in a hanging-drop setup over 500 μL reservoir solution, and the tray was stored at 20°C. Crystals formed within one day and remained stable for the 2.5 weeks between tray setting and crystal freezing. A crystal was looped, submerged in cryoprotection solution (0.05 M sodium succinate (pH 5.3), 0.5 mM spermine, 20 mM magnesium chloride, 3 M ammonium sulfate) for a few seconds, and frozen in liquid nitrogen. Diffraction data were collected under cryogenic conditions at the Advanced Light Source beamline 8.3.1 on a Pilatus3 S 6M (Dectris) detector.

### Data processing, phase determination, and model refinement

Preliminary processing of diffraction images was performed in XDS [43,44]. Unmerged reflections underwent anisotropic truncation, merging, and anisotropic correction using the default parameters of the STARANISO server (v3.339) [45], and a preliminary structural model was included in the input to estimate the expected intensity profile. The best-fit cut-off ellipsoid imposed diffraction limits of 1.656 Å, 2.176 Å, and 1.637 Å based on a cut-off criterion of I/σ(I)=1.2. The “aniso-merged” output MTZ file was used for downstream processing. Using programs within CCP4 (v7.1.015), R_free_ flags were added to 5% of the reflections, and reflections outside the diffraction cut-off surface were removed.

Phases were determined by molecular replacement with Phaser-MR [46], as implemented in Phenix v1.19.2-4158 [47]. The search model comprised two components (unconstrained with respect to each other), both generated in X3DNA v2.4 [48] and each representing one half of the base-paired portion of the crystallization construct. The first component was a 6-base-pair RNA:DNA duplex with perfect A-form geometry and sequence 5’-GCUUAC-3’ / 5’-GTAAGC-3’ (created using the program “fiber” with the -rna option, followed by manual alteration of the DNA strand in PyMOL v2.4.1). The second component was a 6-base-pair DNA:DNA duplex with perfect B-form geometry and sequence 5’-GATGCT-3’ / 5’-AGCATC-3’ (created with “fiber” option −4). Successful phasing was achieved by searching for three copies of each of these components (six components total). Additional phosphodiesters and nucleotides were built in Coot v0.9.2 [49], and the model underwent iterative refinements in Phenix. Phasing and preliminary refinements were initially performed using an earlier (lower-resolution) dataset that had similar unit cell parameters to the final dataset described above.

The initial model, which was refined into a map generated from the earlier dataset, was rigid-body docked into the final-dataset-derived map and underwent further iterative refinements, beginning with resetting of the atomic B-factors, simulated annealing, and addition of ordered solvent. Non-crystallographic symmetry restraints were applied in early rounds of refinement to link the torsion angles of the three molecules within the asymmetric unit; these restraints were removed in the final rounds of refinement. TLSMD [50,51] was used to determine optimal segmentation for Translation/Libration/Screw (TLS) refinement (each 7-mer comprised a separate segment, and the 12-mers were each divided into three segments: nucleotides 1-4, 5-8, 9-12). Refinement using Phenix’s default geometry library yielded dozens of bond lengths and angles that were marked as outliers by the PDB validation server, so the faulty parameters were rigidified *ad hoc* (that is, their estimated standard deviation values in the library files were made smaller, with no change to the mean values). The final three cycles of refinement were performed in Phenix with adjustments to XYZ (reciprocal-space), TLS (segments as indicated above), and individual B-factors. In Table 1, STARANISO and Phenix were used to calculate the data collection statistics and the refinement statistics, respectively.

The final R_free_ value (0.283) is higher than expected for a structure refined using diffraction data at a resolution of 1.6 Å [52]. However, it is important to note that the highest-resolution shell has a completeness of just 6%, and completeness only rises above 95% at ~2.3 Å, due mostly to the anisotropic nature of the diffraction data. Additionally, due to diffraction anisotropy, the 2mF_o_-DF_c_ map appears distorted along certain dimensions, affecting interpretation of Molecule 3 most negatively. Therefore, the geometric details of Molecule 3’s phosphate backbone are poorly constrained, and Molecule 1 or 2 should instead be considered as the most accurate representation of the structure. Anisotropy also prevented identification of water molecules around Molecule 3. Furthermore, the mF_o_-DF_c_ map revealed several globular patches of positive density in the major and minor grooves of all molecules, 3.5-4 Å away from the nearest nucleic acid atom. Because these patches bore no recognizable geometric features, attempts to model them with buffer components failed to improve R_free_, so they were left unmodeled. Any of the mentioned issues may contribute to the high R_free_ value.

Beyond the anisotropy, the overall high B-factors in this structure produce 2mF_o_-DF_c_ density that is “blurred” [53]. To enhance high-resolution features of the map for visual inspection and figure preparation, Coot’s Map Sharpening tool was used. B-factor adjustments used for sharpening are reported in the figure legend. Sharpening only effectively revealed high-resolution features for Molecule 1 or 2, as density from Molecule 3 is too anisotropically distorted.

### Structure analysis and figure preparation

Structural model and map figures were prepared in PyMOL. Alignments were performed using PyMOL’s “align” function without outlier rejection. Regularized A-form and B-form DNA:DNA duplexes were prepared using X3DNA’s “fiber” program (options −1 and −4, respectively), using the same sequence present in the helical portion of the crystallization construct (except RNA was modeled as the corresponding DNA sequence). While the A-form DNA:DNA helix may not perfectly represent a regularized version of the RNA:DNA helix with our sequence [19,20], “fiber” does not permit generation of RNA:DNA helices with generic sequence, and the general geometric features of A-form DNA:DNA vs. A-form RNA:DNA are expected to be similar enough to support the conclusions drawn in this work. Base step and nucleotide geometric parameters were calculated using the “find_pair” and “analyze” programs within X3DNA. On graphs of these parameters, dashed lines indicating the expected value for A-form or B-form DNA were calculated by performing an equivalent analysis on the X3DNA-generated regularized A-form/B-form helices and taking the average across all base steps/nucleotides, unless indicated otherwise. Nucleotides with A/B character exhibit a spread of values around those indicated by the dashed lines (as represented more accurately by the dashed ellipses in Fig 4B), and the dashed lines are drawn merely to guide the reader’s eye to general trends. Angles between the helical axes of the

DNA:DNA and RNA:DNA duplex were calculated as the angle between the helical axis vectors of the aligned regularized A-form and B-form helices. Graphs were prepared using matplotlib v3.3.2 [54]. Final figures were prepared in Adobe Illustrator v25.4.1.

## Acknowledgements

We thank J.M. Holton and J.H. Cate for data processing advice. We thank G. Meigs for technical assistance at the beamline. We thank J. Kuriyan for scientific advice.

## Supporting information

**S1 Fig. Additional geometric details of the A-to-B transition.**

(A) X-displacement of the 11 base steps of the 12-bp helix. Black, DNA; red, RNA. (B) Inclination of the 11 base steps of the 12-bp helix. (C) Slide of the 11 base steps of the 12-bp helix. (D) Pseudorotation phase angles for the ribose/deoxyribose conformation at every nucleotide within the 12-bp helix (24 data points per molecule). The modeled sugar conformations might not be unique solutions for this dataset, as in many cases these structural details cannot be directly discerned from the 2mF_o_-DF_c_ map. For this dataset, the most reliable parameters are those defined directly by the nucleobase and phosphate positions, which appear clearly in the 2mF_o_-DF_c_ map (and likely impose indirect geometric constraints on the sugar pucker).

S2 Fig.

**S2 Fig. Nucleotide torsion angles for Molecules 2 and 3.**

Analogous to Fig 4B.

## References

1. Broadwater DWB, Cook AW, Kim HD. First passage time study of DNA strand displacement. Biophys J. 2021;120: 2400–2412. doi:10.1016/j.bpj.2021.01.043

2. Hays FA, Watson J, Ho PS. Caution! DNA crossing: crystal structures of Holliday junctions. J Biol Chem. 2003;278: 49663–49666. doi:10.1074/jbc.R300033200

3. Kowalczykowski SC. Biochemistry of genetic recombination: energetics and mechanism of DNA strand exchange. Annu Rev Biophys Biophys Chem. 1991;20: 539–575. doi: 10.1146/annurev.bb.20.060191.002543

4. McKinney SA, Déclais A-C, Lilley DMJ, Ha T. Structural dynamics of individual Holliday junctions. Nat Struct Biol. 2003;10: 93–97. doi:10.1038/nsb883

5. Ortiz-Lombardía M, González A, Eritja R, Aymamí J, Azorín F, Coll M. Crystal structure of a DNA Holliday junction. Nat Struct Biol. 1999;6: 913–917. doi:10.1038/13277

6. Seeman NC, Kallenbach NR. DNA branched junctions. Annu Rev Biophys Biomol Struct. 1994;23: 53–86. doi:10.1146/annurev.bb.23.060194.000413

7. Simmel FC, Yurke B, Singh HR. Principles and Applications of Nucleic Acid Strand Displacement Reactions. Chem Rev. 2019;119: 6326–6369. doi:10.1021/acs.chemrev.8b00580

8. Srinivas N, Ouldridge TE, Sulc P, Schaeffer JM, Yurke B, Louis AA, et al. On the biophysics and kinetics of toehold-mediated DNA strand displacement. Nucleic Acids Res. 2013;41: 10641–10658. doi:10.1093/nar/gkt801

9. Crossley MP, Bocek M, Cimprich KA. R-Loops as Cellular Regulators and Genomic Threats. Mol Cell. 2019;73: 398–411. doi:10.1016/j.molcel.2019.01.024

10. Altae-Tran H, Kannan S, Demircioglu FE, Oshiro R, Nety SP, McKay LJ, et al. The widespread IS200/IS605 transposon family encodes diverse programmable RNA-guided endonucleases. Science. 2021;374: 57–65. doi:10.1126/science.abj6856

11. Karvelis T, Druteika G, Bigelyte G, Budre K, Zedaveinyte R, Silanskas A, et al. Transposon-associated TnpB is a programmable RNA-guided DNA endonuclease. Nature. 2021;599: 692–696. doi:10.1038/s41586-021-04058-1

12. Chen JS, Doudna JA. The chemistry of Cas9 and its CRISPR colleagues. Nat Rev Chem. 2017;1: 1–15. doi:10.1038/s41570-017-0078

13. Knott GJ, Doudna JA. CRISPR-Cas guides the future of genetic engineering. Science. 2018;361: 866–869. doi:10.1126/science.aat5011

14. Pickar-Oliver A, Gersbach CA. The next generation of CRISPR-Cas technologies and applications. Nat Rev Mol Cell Biol. 2019;20: 490–507. doi:10.1038/s41580-019-0131-5

15. Sternberg SH, Redding S, Jinek M, Greene EC, Doudna JA. DNA interrogation by the CRISPR RNA-guided endonuclease Cas9. Nature. 2014;507: 62–67. doi:10.1038/nature13011

16. Cofsky JC, Karandur D, Huang CJ, Witte IP, Kuriyan J, Doudna JA. CRISPR-Cas12a exploits R-loop asymmetry to form double-strand breaks. Wolberger C, Bailey S, Ke A, White MF, editors. eLife. 2020;9: e55143. doi:10.7554/eLife.55143

17. Aymami J, Coll M, Marel GA van der, Boom JH van, Wang AH, Rich A. Molecular structure of nicked DNA: a substrate for DNA repair enzymes. PNAS. 1990;87: 2526–2530.

18. Roll C, Ketterlé C, Faibis V, Fazakerley GV, Boulard Y. Conformations of nicked and gapped DNA structures by NMR and molecular dynamic simulations in water. Biochemistry. 1998;37: 4059–4070. doi:10.1021/bi972377w

19. Fedoroff OYu null, Salazar M, Reid BR. Structure of a DNA:RNA hybrid duplex. Why RNase H does not cleave pure RNA. J Mol Biol. 1993;233: 509–523. doi:10.1006/jmbi.1993.1528

20. Horton NC, Finzel BC. The Structure of an RNA/DNA Hybrid: A Substrate of the Ribonuclease Activity of HIV-1 Reverse Transcriptase. Journal of Molecular Biology. 1996;264: 521–533. doi:10.1006/jmbi.1996.0658

21. Milman G, Langridge R, Chamberlin MJ. The structure of a DNA-RNA hybrid. Proc Natl Acad Sci U S A. 1967;57: 1804–1810.

22. Zhu L, Salazar M, Reid BR. DNA duplexes flanked by hybrid duplexes: the solution structure of chimeric junctions in [r(cgcg)d(TATACGCG)]2. Biochemistry. 1995;34: 2372–2380. doi:10.1021/bi00007a033

23. Dickerson RE, Goodsell DS, Neidle S. …the tyranny of the lattice… Proc Natl Acad Sci U S A. 1994;91: 3579–3583. doi:10.1073/pnas.91.9.3579

24. Drew HR, Wing RM, Takano T, Broka C, Tanaka S, Itakura K, et al. Structure of a B-DNA dodecamer: conformation and dynamics. Proc Natl Acad Sci U S A. 1981;78: 2179–2183. doi:10.1073/pnas.78.4.2179

25. Saenger W. Principles of Nucleic Acid Structure. Springer Science & Business Media; 1984.

26. El Hassan MA, Calladine CR. Conformational characteristics of DNA: empirical classifications and a hypothesis for the conformational behaviour of dinucleotide steps. Philosophical Transactions of the Royal Society of London Series A: Mathematical, Physical and Engineering Sciences. 1997;355: 43–100. doi:10.1098/rsta.1997.0002

27. Lu XJ, Shakked Z, Olson WK. A-form conformational motifs in ligand-bound DNA structures. J Mol Biol. 2000;300: 819–840. doi:10.1006/jmbi.2000.3690

28. Ban C, Ramakrishnan B, Sundaralingam M. A single 2’-hydroxyl group converts B-DNA to A-DNA. Crystal structure of the DNA-RNA chimeric decamer duplex d(CCGGC)r(G)d(CCGG) with a novel intermolecular G-C base-paired quadruplet. J Mol Biol. 1994;236: 275–285. doi:10.1006/jmbi.1994.1134

29. Egli M, Usman N, Zhang SG, Rich A. Crystal structure of an Okazaki fragment at 2-A resolution. Proc Natl Acad Sci U S A. 1992;89: 534–538. doi:10.1073/pnas.89.2.534

30. Egli M, Usman N, Rich A. Conformational influence of the ribose 2’-hydroxyl group: crystal structures of DNA-RNA chimeric duplexes. Biochemistry. 1993;32: 3221–3237.

31. Wahl MC, Sundaralingam M. B-form to A-form conversion by a 3’-terminal ribose: crystal structure of the chimera d(CCACTAGTG)r(G). Nucleic Acids Res. 2000;28: 4356–4363. doi:10.1093/nar/28.21.4356

32. Wang AH, Fujii S, van Boom JH, van der Marel GA, van Boeckel SA, Rich A. Molecular structure of r(GCG)d(TATACGC): a DNA--RNA hybrid helix joined to double helical DNA. Nature. 1982;299: 601–604. doi:10.1038/299601a0

33. Mellema JR, Haasnoot CA, van der Marel GA, Wille G, van Boeckel CA, van Boom JH, et al. Proton NMR studies on the covalently linked RNA-DNA hybrid r(GCG)d(TATACGC). Assignment of proton resonances by application of the nuclear Overhauser effect. Nucleic Acids Res. 1983;11: 5717–5738. doi:10.1093/nar/11.16.5717

34. Salazar M, Fedoroff OYu null, Zhu L, Reid BR. The solution structure of the r(gcg)d(TATACCC):d(GGGTATACGC) Okazaki fragment contains two distinct duplex morphologies connected by a junction. J Mol Biol. 1994;241: 440–455. doi:10.1006/jmbi.1994.1519

35. Salazar M, Fedoroff OY, Reid BR. Structure of chimeric duplex junctions: solution conformation of the retroviral Okazaki-like fragment r(ccca)d(AATGA).d(TCATTTGGG) from Moloney murine leukemia virus. Biochemistry. 1996;35: 8126–8135. doi:10.1021/bi9528917

36. Selsing E, Wells RD, Early TA, Kearns DR. Two contiguous conformations in a nucleic acid duplex. Nature. 1978;275: 249–250. doi:10.1038/275249a0

37. Häse F, Zacharias M. Free energy analysis and mechanism of base pair stacking in nicked DNA. Nucleic Acids Res. 2016;44: 7100–7108. doi:10.1093/nar/gkw607

38. Pacesa M, Jinek M. Mechanism of R-loop formation and conformational activation of Cas9. 2021. p. 2021.09.16.460614. Available: https://www.biorxiv.org/content/10.1101/2021.09.16.460614v1

39. Zhu X, Clarke R, Puppala AK, Chittori S, Merk A, Merrill BJ, et al. Cryo-EM structures reveal coordinated domain motions that govern DNA cleavage by Cas9. Nat Struct Mol Biol. 2019;26: 679–685. doi:10.1038/s41594-019-0258-2

40. Cofsky JC, Soczek KM, Knott GJ, Nogales E, Doudna JA. CRISPR-Cas9 bends and twists DNA to read its sequence. Biochemistry; 2021 Sep. doi:10.1101/2021.09.06.459219

41. Lapinaite A, Knott GJ, Palumbo CM, Lin-Shiao E, Richter MF, Zhao KT, et al. DNA capture by a CRISPR-Cas9-guided adenine base editor. Science. 2020;369: 566–571. doi:10.1126/science.abb1390

42. Cavaluzzi MJ, Borer PN. Revised UV extinction coefficients for nucleoside-5’-monophosphates and unpaired DNA and RNA. Nucleic Acids Res. 2004;32: e13. doi:10.1093/nar/gnh015

43. Kabsch W. XDS. Acta Cryst D. 2010;66: 125–132. doi:10.1107/S0907444909047337

44. Kabsch W. Integration, scaling, space-group assignment and post-refinement. Acta Cryst D. 2010;66: 133–144. doi:10.1107/S0907444909047374

45. Tickle IJ, Flensburg C, Keller P, Paciorek W, Sharff A, Vonrhein C, et al. STARANISO. Cambridge, United Kingdom: Global Phasing Ltd.; 2018. Available: http://staraniso.globalphasing.org/cgi-bin/staraniso.cgi

46. McCoy AJ, Grosse-Kunstleve RW, Adams PD, Winn MD, Storoni LC, Read RJ. Phaser crystallographic software. J Appl Cryst. 2007;40: 658–674. doi:10.1107/S0021889807021206

47. Liebschner D, Afonine PV, Baker ML, Bunkóczi G, Chen VB, Croll TI, et al. Macromolecular structure determination using X-rays, neutrons and electrons: recent developments in Phenix. Acta Cryst D. 2019;75: 861–877. doi:10.1107/S2059798319011471

48. Lu X-J, Olson WK. 3DNA: a software package for the analysis, rebuilding and visualization of three-dimensional nucleic acid structures. Nucleic Acids Res. 2003;31: 5108–5121. doi:10.1093/nar/gkg680

49. Emsley P, Lohkamp B, Scott WG, Cowtan K. Features and development of Coot. Acta Crystallogr D Biol Crystallogr. 2010;66: 486–501. doi:10.1107/S0907444910007493

50. Painter J, Merritt EA. TLSMD web server for the generation of multi-group TLS models. J Appl Cryst. 2006;39: 109–111. doi:10.1107/S0021889805038987

51. Painter J, Merritt EA. Optimal description of a protein structure in terms of multiple groups undergoing TLS motion. Acta Cryst D. 2006;62: 439–450. doi:10.1107/S0907444906005270

52. Kleywegt GJ, Brünger AT. Checking your imagination: applications of the free R value. Structure. 1996;4: 897–904. doi:10.1016/s0969-2126(96)00097-4

53. Liu C, Xiong Y. Electron density sharpening as a general technique in crystallographic studies. J Mol Biol. 2014;426: 980–993. doi:10.1016/j.jmb.2013.11.014

54. Hunter JD. Matplotlib: A 2D Graphics Environment. Computing in Science Engineering. 2007;9: 90–95. doi:10.1109/MCSE.2007.55

